# Dorsal color variation among subspecies of the “Oregon” Dark-eyed Junco complex

**DOI:** 10.1101/2021.01.24.427985

**Authors:** Elisa T. Yang, Philip Unitt, Nicholas A. Mason

## Abstract

Initial descriptions of avian subspecies were based on expert opinions of geographic variation in phenotypes and are inherently subjective. Although best practices for subspecies delimitation continue to be debated, reassessing subspecies limits with current, quantitative methods is important toward refining and improving taxonomic treatments. Plumage coloration is the basis of many subspecies diagnoses, but is potentially problematic because of the historical lack of quantitative methods to quantify color. Recently developed methods, such as colorimetry, provide repeatable measurements of color variation that can be used to reassess subspecies limits. In this study, we reassess color variation among subspecies of the Oregon Dark-eyed Junco (*Junco hyemalis [oreganus Group]*) complex, in which differences in back and hood color were established as diagnostic characters. We measured back and hood brightness and hue in 206 museum specimens among five Oregon Dark-eyed Junco subspecies using a colorimeter. We then compared mean measurements among subspecies and conducted a discriminant function analysis to assess how well dorsal color predicted subspecies. Our data correctly classified only 67.9% of males and 82.5% of females to their presumed subspecies. Furthermore, no adjacent subspecies pairs passed the “75% rule” due to extensive overlap in plumage characters. Thus, back color alone is not as effective in diagnosing Oregon Dark-eyed Junco subspecies as originally described, suggesting a possible taxonomic revision. Specifically, similarity in phenotypic and genetic data suggests that some combination of *thurberi*, *montanus*, and *shufeldti* may be lumped to recognize broad, clinal variation in dorsal color alongside clinal variation in other phenotypes and extensive gene flow.

---

Avian subspecies delimitation is a controversial taxonomic practice that has been debated and refined since its initial conceptualization (Mayr 1943; Wilson & Brown 1953; Zink 2004; Patten 2010, 2015; Remsen 2010; Winker 2010). In birds, subspecies are generally thought to represent phenotypically and/or genotypically differentiated populations within a species that occupy a geographic region (Patten & Remsen 2017). However, original subspecies descriptions were based mostly on expert opinions of geographic variation in phenotypes, resulting in subjective classifications that often fail to meet current best practices (James 2010). Today, delimiting subspecies is still far from straightforward: taxonomists continue to differ on what subspecies represent and optimal criteria for delimitation (Fitzpatrick 2010; Gill 2014; Cronin et al. 2015; Weckworth et al. 2015; Vinarski 2015). Nonetheless, the development and implementation of quantitative metrics and statistical analyses promote a more consistent and standardized subspecies classification system (Patten 2010). Many subspecies described prior to the development of current practices are equivocal and would benefit from reassessments using modern approaches. Such reassessments are important for current applied and basic research initiatives, many of which use subspecies as units of biodiversity and indices of ecological and evolutionary processes (Phillimore & Owens 2006; Haig & D’Elia 2010).

Phenotypic variation guided initial subspecies descriptions and remains important in infraspecific taxonomy, even as genetic data play a growing role in subspecies delimitation (Winker 2009; Patten & Remsen 2017). Coloration in particular has played a prominent role in avian taxonomy for multiple reasons. First, plumage coloration and patterning is influenced by selective pressures, such as natural selection favoring coloration that promotes camouflage or sexual selection favoring bright, showy colors that promote mate choice (Hill & McGraw 2006; Mason & Bowie 2020). Thus, differences among populations may represent evolutionary changes in response to local conditions (Zink & Remsen 1986; Zamudio et al. 2016). Second, differences in coloration are readily observable by the human eye and were easily detected by early taxonomists (Endler 1990). However, historical assessments of color differences relied on individual assessment of matches between plumage patches and color swatches (Ridgway 1912), which may introduce qualitative, subjective differences among observers (Zuk & Decruyenaere 1994; Butler et al. 2011). Today, colorimetry, spectrophotometry, and digital photography offer affordable ways to accurately measure color variation in a consistent, quantitative manner (Burns et al. 2017). Recently, colorimetry has been used to quantify color variation and reexamine subspecific taxonomy in various groups, including Willow Flycatchers (*Empidonax trailli*; Paxton et al. 2010), Least Terns (*Sternula antillarum*; Johnson et al. 1998), and Sagebrush / Bell’s Sparrows (*Artemisiospiza sp.*; Patten & Unitt 2002). Nonetheless, many subspecies groups are still in need of quantitative reevaluations of color variation and diagnosability among taxa.

Among the subspecies groups that would benefit from a reexamination of how coloration differences correspond to subspecies is the Oregon Dark-eyed Junco (*Junco hyemalis*) complex. Dark-eyed Juncos exhibit pronounced intraspecific plumage variation, with seven ‘groups’ of subspecies that have recently diversified across North America (Milá et al. 2007; Friis et al. 2016; Clements et al. 2019). In western North America, from Baja California north to Alaska, seven subspecies comprise the “Oregon” Dark-eyed Junco group (Clements et al. 2019; Nolan et al. 2020). Following comprehensive analyses of Oregon Junco subspecies by early taxonomists (Ridgway 1901; Dwight 1918), Miller (1941) further established the current taxonomy of the genus *Junco* by matching the specimens’ hoods and backs with graded color samples and examining pigments under a microscope (see Table 1 for subspecies descriptions). Despite the widespread, longstanding use of Miller’s (1941) classification, it is still unknown whether these subspecies represent diagnosable taxa that meet current guidelines for subspecies delimitation.

**Table 1:**
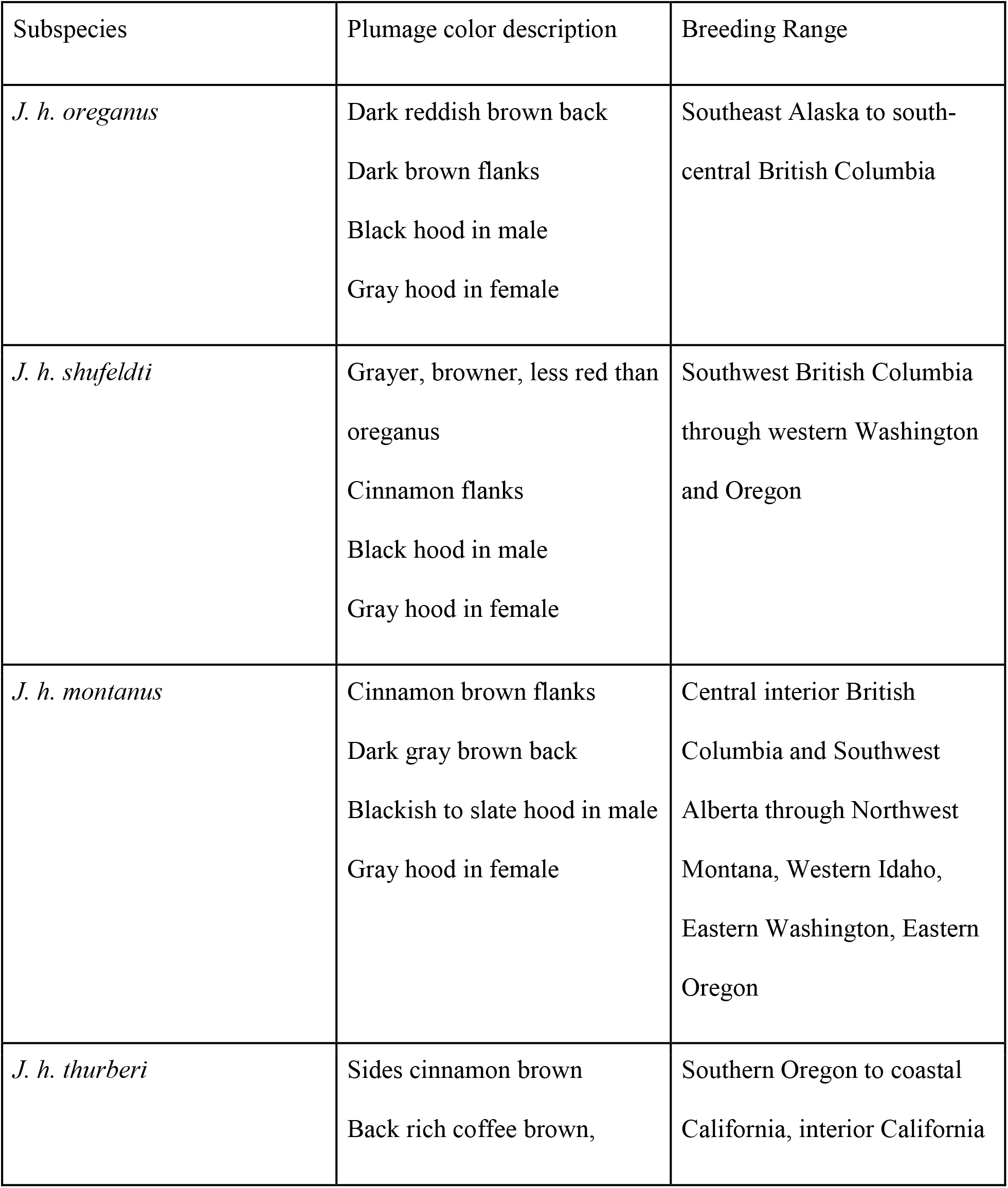
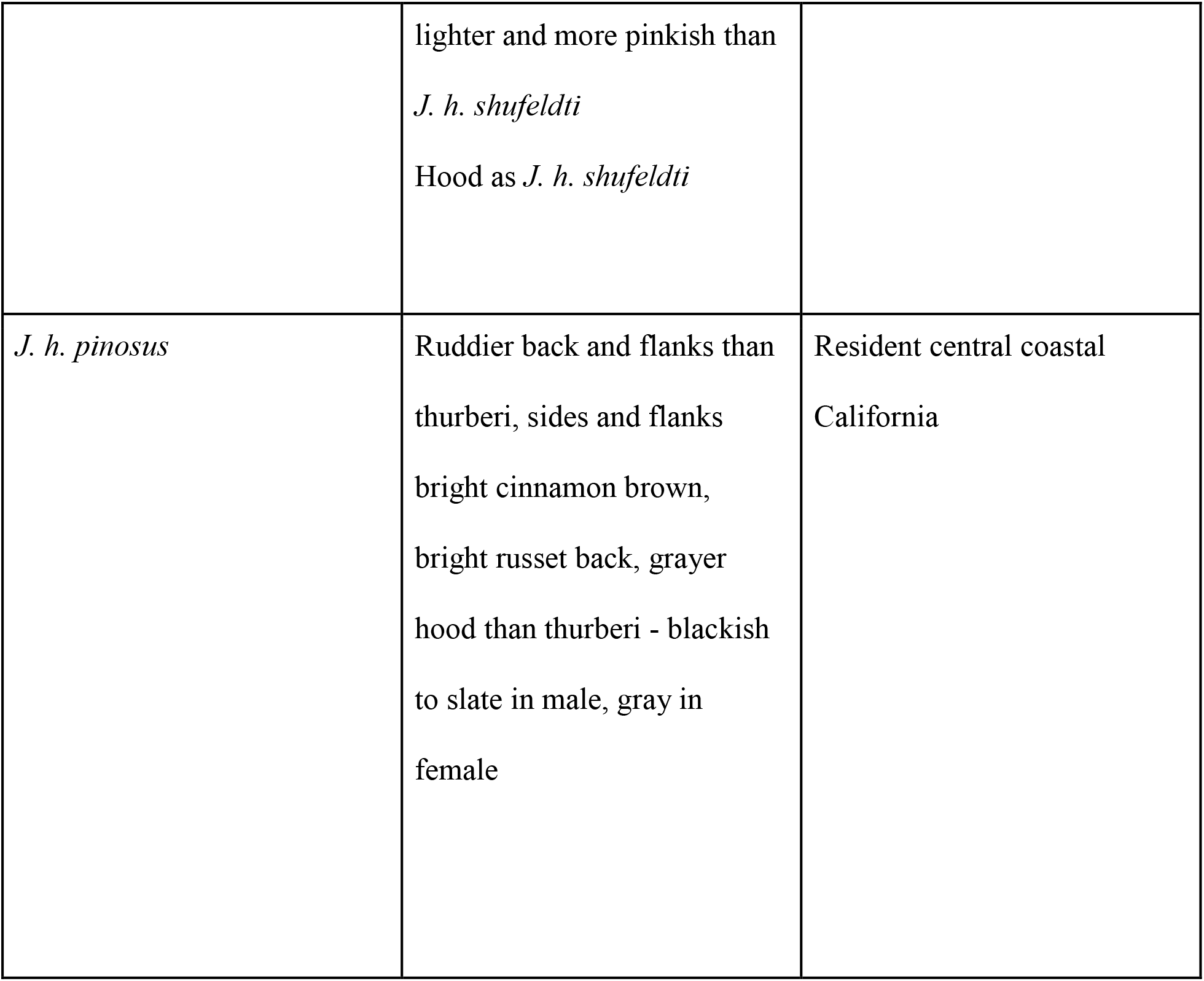
Phenotypic descriptions of the five subspecies’ appearances and breeding ranges (following Miller (1941)) included in this study.

In this study, we reevaluated Miller’s (1941) classification of the Oregon Dark-eyed Junco complex using colorimetry. We compared back and hood color variation between five subspecies from the Oregon Dark-eyed Junco complex (*pinosus*, *thurberi*, *shufeldti*, *montanus*, and *oreganus*), and excluded two subspecies from Mexico (*pontillis*, *townsendi*) for which we lacked adequate sample sizes. We compared mean values of brightness and hue measurements among sexes, age classes, and subspecies. We also quantified subspecies diagnosability of males and females using a discriminant function analysis. Finally, we tested the “75% rule” (Amadon 1949; Patten & Unitt 2002) to see if quantitative color variation among subspecies passed a widely used ‘yardstick’ of diagnosability. In doing so, we reassessed the validity of back color as a diagnostic character for Oregon Dark-eyed Junco subspecies, and reconsidered subspecies limits within the complex.

## Methods

We measured plumage reflectance from 206 specimens of Oregon Dark-eyed Junco (Figure 1; Appendix A). Our sampling was drawn from specimens housed in three collections: the San Diego Natural History Museum (SDNHM), Natural History Museum of Los Angeles County, Los Angeles (LACM), and the University of Los Angeles, California (UCLA). We measured a minimum of 15 individuals from 5 subspecies, including (from north to south) *J. h. oreganus* (16), *J. h. montanus* (29), *J. h. shufeldti* (17), *J. h. thurberi* (125), *J. h. pinosus* (n = 19). Specimens were selected based on the ranges described by Miller. Individuals falling out of the range of their labeled subspecies were removed. Wintering individuals in subspecies with nonbreeding ranges overlapping the range of other subspecies were also removed.

**Figure 1:**
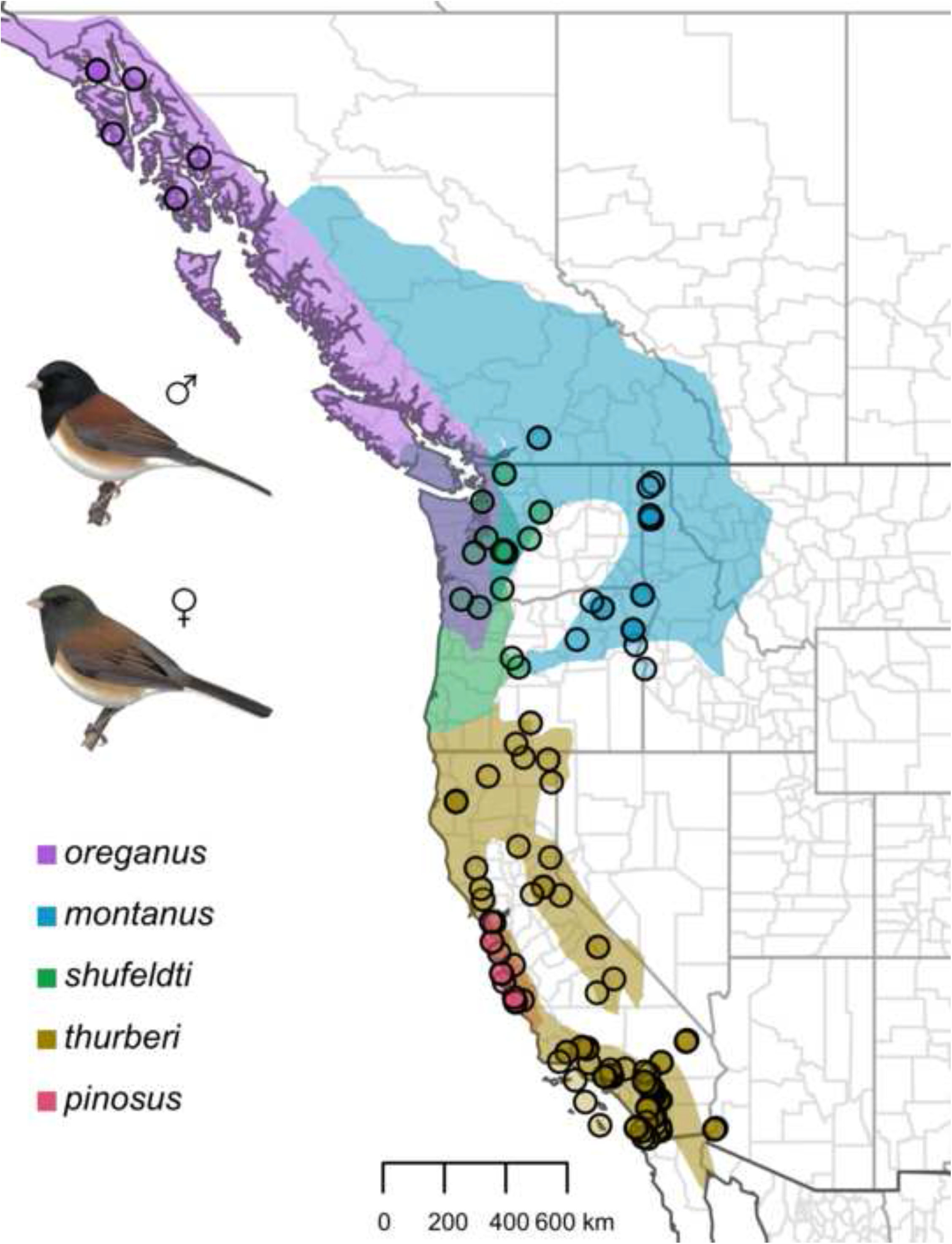
Sampling map of vouchered specimens included in our study of plumage reflectance among subspecies of the “Oregon” Dark-eyed Junco. Plates of male and female “Oregon” Dark-eyed Junco are shown on the left. Approximate breeding ranges of each subspecies are shown in different colors and have been modified following the descriptions of Miller (1941). As our study included wintering as well as breeding birds, some circles fall outside of their expected range due to non-breeding seasonal movements. Sampling localities are indicated with circles with subspecies indicated by the fill color of the circle. Some dots may represent more than one individual sampled from the same locality. Illustrations are provided courtesy of Lynx Edicions.

We did not include specimens with significantly worn plumage nor those that had faded or missing feathers. We also omitted juveniles with irregular streaking, diffuse brown dorsal plumage, obvious truncations in tail or greater coverts or completely unossified skulls. Immature birds were identified whenever possible via known aging criteria, tag labels noting partially ossified skulls, or labels noting developing ovaries. Birds for which none of this information was obvious or available were labeled as adult. Thus, each individual included in the study was classified as either immature or adult.

We measured the coloration of the center of the hood and the back of each specimen using a Konica Minolta CR-300 Chroma Meter (Ramsey, New Jersey, USA). We recorded three measurements: (1) *L*\*, or brightness, in which higher values corresponding to brighter plumage; (2) *a*\*, or redness, in which lower values correspond to greener hues and higher values correspond redder hues; and (3) *b*\*, or yellowness, in which lower values correspond to bluer hues and higher values correspond to yellower hues. We repeated each measurement three times and subsequently averaged those measurements.

We constructed separate generalized linear models (GLMs) for each of the six dorsal color measurements (hood *L*, hood *a*, hood *b*, back *L*, back *a*, and back *b*) with a Gaussian distribution of error using the glm() function in the R programming environment (R Core Team 2020). We included subspecies, sex, age class, days since molt, and years since collection as main effects in each model. Dark-eyed Juncos undergo two body molts per year: an alternate plumage molt from February to April and a basic molt from July to October (Pyle 1997).

Definitive basic and alternate plumages are nearly identical in Dark-eyed Juncos (Nolan et al. 2020), but color may change as freshly molted feathers wear and abrade over time (Tökölyi et al. 2008). We therefore incorporated the number of days since molt into our GLMs as the difference between the collection date and the most recent molt event of either March 15th or September 1st. Furthermore, because specimens can fade and change color over years since their initial collection (Doucet & Hill 2009), we also incorporated the number of years since collection into our GLMs. We examined the distribution of residuals for each model to ensure that they approximated a normal distribution (Supplementary Figure SF1). We subsequently performed a series of Tukey’s Honestly Significant Difference (HSD) tests to quantify differences in the mean values of each plumage metric among sexes, age classes, and subspecies using the R package agricolae v1.3.1 (de Mendiburu 2019).

We also performed a discriminant function analysis (DFA) on adult males (n=91) and females (n=83) separately to determine the diagnosability of Oregon Junco subspecies based on hood and mantle coloration using the MASS package (Venables & Ripley 2002) in R. We subsequently performed a ‘leave-one-out’ (i.e., jackknifed) cross validation on our DFA to predict the subspecies of each individual specimen and test the diagnosability of each subspecies for both sexes (i.e., quantify the proportion of individuals that were correctly predicted as their identified subspecies based on colorimetry data).

Finally, we implemented a widely-used diagnosability test commonly referred to as the “75% rule” (Amadon 1949; Patten & Unitt 2002). In brief, the “75% rules” involves a pair-wise test that determines whether 75% of the distribution of a single trait for one subspecies falls outside of 99% of another subspecies’ distribution for the same trait. The derivation of the test statistic is described in detail in Patten and Unitt (2002) and is shown here.

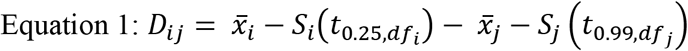

In this equation, 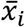 the mean and *S*_*j*_ is the standard deviation of a given trait for subspecies *i*. If the *D*_*ij*_ statistic is greater than 1, then subspecies *i* is diagnosable from subspecies *j*. For the “75% rule” to be met, the converse has to be true such that both subspecies are diagnosable from each other. The second statistic (*D*_*ji*_) can be calculated by swapping the critical *t* values in the equation above. In our case, we tested pairwise diagnosability via the “75% rule” for each subspecies pair with overlapping ranges. We only included adults and considered each sex separately. This resulted in five diagnosability tests (from northernmost to southernmost): (1) *oreganus* and *montanus*, (2) *oreganus* and *shufeldti*, (3) *montanus* and *shufeldti*, (4) *shufeldti* and *thurberi*, and (5) *thurberi* and *pinosus*. We conducted a DFA for each subspecies pair and then used scores from the first DFA axis to calculate the two summary statistics, *D*_*ij*_ and *D*_*ji*_ for adult males and females.

## Results

We uncovered variation in hood and mantle coloration among sexes, age classes, and five subspecies within the “Oregon” Dark-eyed Junco complex. When we compared hood and back color among males and females, we found that females had higher hood *L* and hood *b* values, but lower hood *a* values (Fig. 2, Table 2). Among back measurements, females had higher back *L*, back *a*, and back *b* values compared to males (Fig. 2, Table 2). Overall, this indicates that females typically have lighter hoods and backs compared to males, while female hoods and backs are typically ‘buffier’ with higher red and yellow reflectance than males. Although average values differed between males and females, there was still substantial overlap in the range of plumage coloration measurements between sexes.

**Table 2:** Results from generalized linear model (GLM) analyses for each of six response variables: hood *L*, hood *a*, hood *b*, back *L*, back *a*, back *b*. Significant model effects (P < 0.05) are displayed with a bold font for the P value. GLMs were generated with a Gaussian distribution of error. Categorical terms are compared to a base model with male sex, adult age class, and *oreganus* subspecies. Days since molt Specimen age displays change in parameter values per year.

| Hood L | $\beta \pm SE$ | t value | P value |
| --- | --- | --- | --- |
| (Intercept) | $27.183 \pm 0.617$ | 44.036 | **>0.0001** |
| Sex Female | $0.768 \pm 0.267$ | 2.875 | **0.0045** |
| Age Class Immature | $-0.772 \pm 0.314$ | -2.463 | **0.0146** |
| Subspecies *montanus* | $4.118 \pm 0.589$ | 6.991 | **>0.0001** |
| Subspecies *shufeldti* | $3.4 \pm 0.641$ | 5.302 | **>0.0001** |
| Subspecies *thurberi* | $5.861 \pm 0.53$ | 11.05 | **>0.0001** |
| Subspecies *pinosus* | $3.826 \pm 0.627$ | 6.103 | **>0.0001** |
| Days Since Molt | $0.002 \pm 0.003$ | 0.798 | 0.4261 |
| Specimen Age | $-0.004 \pm 0.004$ | -1.012 | 0.3126 |
| Hood a | $\beta \pm SE$ | t value | P value |
| (Intercept) | $5.156 \pm 0.259$ | 19.928 | **>0.0001** |
| Sex Female | $-0.243 \pm 0.112$ | -2.168 | **0.0313** |
| Age Class Immature | $0.458 \pm 0.131$ | 3.485 | **0.0006** |
| Subspecies *montanus* | $-1.401 \pm 0.247$ | -5.676 | **>0.0001** |
| Subspecies *shufeldti* | $-0.683 \pm 0.269$ | -2.543 | **0.0117** |
| Subspecies *thurberi* | $0.014 \pm 0.222$ | 0.065 | 0.9483 |
| Subspecies *pinosus* | $1.555 \pm 0.263$ | 5.92 | **>0.0001** |
| Days Since Molt | $-0.001 \pm 0.001$ | -0.974 | 0.3312 |
| Specimen Age | $0.002 \pm 0.002$ | 1.075 | 0.2837 |

| Hood b | $\beta \pm SE$ | t value | P value |
| --- | --- | --- | --- |
| (Intercept) | $13.171 \pm 0.436$ | 30.195 | **>0.0001** |
| Sex Female | $0.542 \pm 0.189$ | 2.869 | **0.0046** |
| Age Class Immature | $0.194 \pm 0.222$ | 0.874 | 0.383 |
| Subspecies *montanus* | $-1.867 \pm 0.416$ | -4.485 | **>0.0001** |
| Subspecies *shufeldti* | $-0.333 \pm 0.453$ | -0.734 | 0.4639 |
| Subspecies *thurberi* | $1.123 \pm 0.375$ | 2.996 | **0.0031** |
| Subspecies *pinosus* | $2.466 \pm 0.443$ | 5.566 | **>0.0001** |
| Days Since Molt | $-0.002 \pm 0.002$ | -1.047 | 0.2966 |
| Specimen Age | $0.016 \pm 0.003$ | 6.128 | **>0.0001** |
| Back L | $\beta \pm SE$ | t value | P value |
| (Intercept) | $15.892 \pm 1.102$ | 14.416 | **>0.0001** |
| Sex Female | 5.988 ± 0.477 | 12.555 | **>0.0001** |
| Age Class Immature | 2.839 ± 0.56 | 5.068 | **>0.0001** |
| Subspecies *montanus* | 2.112 ± 1.052 | 2.007 | **0.0461** |
| Subspecies *shufeldti* | 1.207 ± 1.145 | 1.054 | 0.2933 |
| Subspecies *thurberi* | 1.242 ± 0.947 | 1.312 | 0.1912 |
| Subspecies *pinosus* | 1.548 ± 1.12 | 1.383 | 0.1682 |
| Days Since Molt | 0.006 ± 0.005 | 1.261 | 0.2087 |
| Specimen Age | 0.007 ± 0.007 | 1.082 | 0.2804 |

| Back a | $\beta \pm SE$ | t value | P value |
| --- | --- | --- | --- |
| (Intercept) | 1.296 ± 0.203 | 6.396 | **>0.0001** |
| Sex Female | 0.417 ± 0.088 | 4.757 | **>0.0001** |
| Age Class Immature | 0.627 ± 0.103 | 6.089 | **>0.0001** |
| Subspecies *montanus* | -0.187 ± 0.193 | -0.967 | 0.3348 |
| Subspecies *shufeldti* | 0.154 ± 0.211 | 0.732 | 0.4647 |
| Subspecies *thurberi* | 0.158 ± 0.174 | 0.907 | 0.3657 |
| Subspecies *pinosus* | 0.537 ± 0.206 | 2.609 | **0.0098** |
| Days Since Molt | -0.001 ± 0.001 | -1.651 | 0.1004 |
| Specimen Age | 0.008 ± 0.001 | 6.251 | **>0.0001** |
| Back b | $\beta \pm SE$ | t value | P value |
| (Intercept) | 4.053 ± 0.582 | 6.959 | **>0.0001** |
| Sex Female | 2.196 ± 0.252 | 8.713 | **>0.0001** |
| Age Class Imature | 2.11 ± 0.296 | 7.132 | **>0.0001** |
| Subspecies *montanus* | -0.768 ± 0.556 | -1.382 | 0.1687 |
| Subspecies *shufeldti* | 0.375 ± 0.605 | 0.619 | 0.5364 |
| Subspecies *thurberi* | 0.32 ± 0.5 | 0.64 | 0.5231 |
| Subspecies *pinosus* | 1.29 ± 0.591 | 2.182 | 0.0303 |
| Days Since Molt | -0.003 ± 0.003 | -1.366 | 0.1736 |
| Specimen Age | 0.019 ± 0.003 | 5.488 | **>0.0001** |

**Figure 2:**
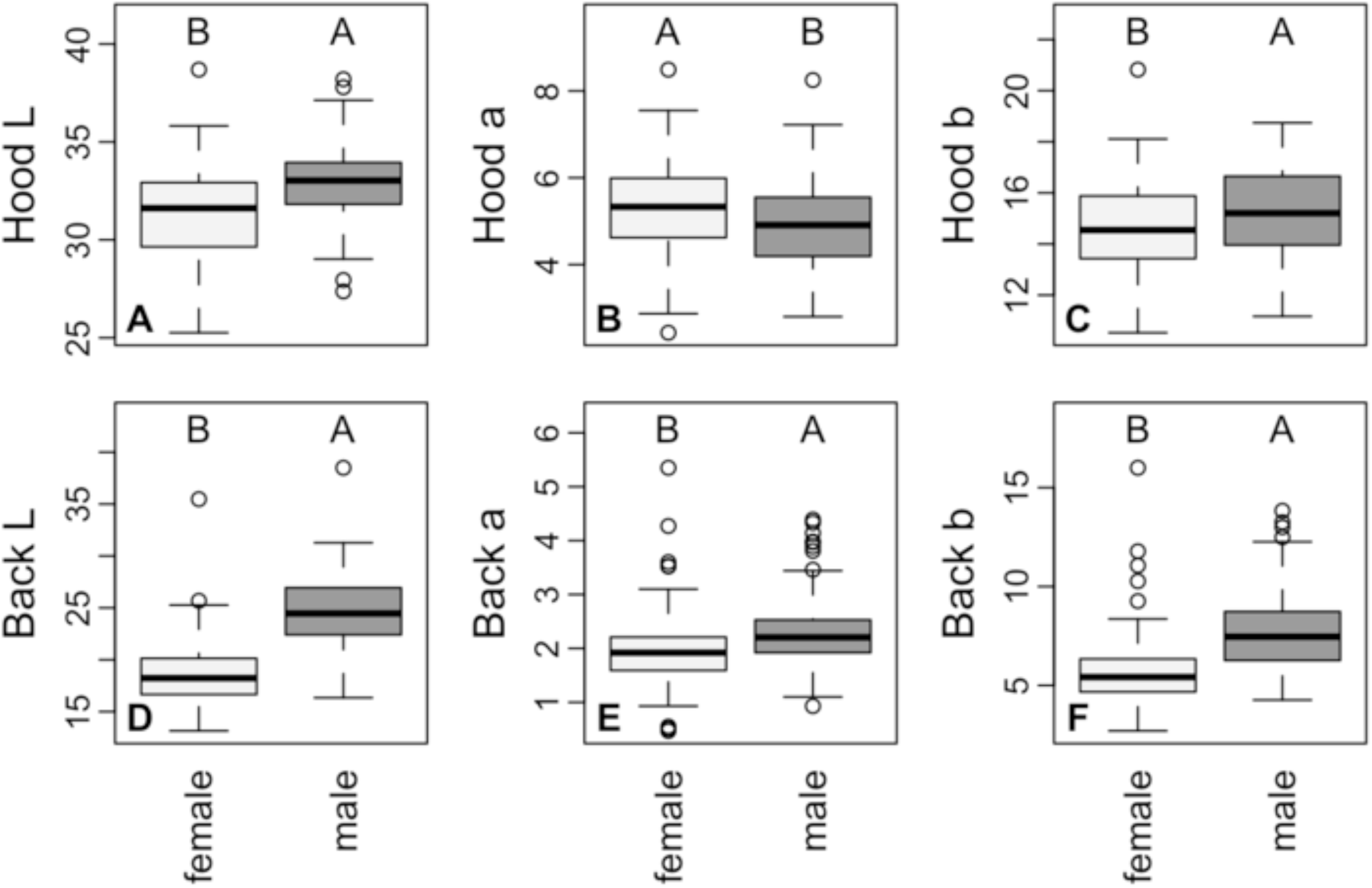
Boxplots of hood and back coloration measurements among males and females of the “Oregon” Dark-eyed Junco complex. For the back and hood, “L” corresponds to brightness with higher values indicating brighter colors; “a” corresponds to redness, in which higher values correspond to more red coloration; and “b” corresponds to yellowness, in which higher values correspond to more yellow coloration. Shown above each box is the group classification following a Tukey’s HSD test for pairwise differences in mean values with the alphabetical order of groupings corresponding to descending differences in mean values among groups.

Age classes also differed in dorsal color. Hood *a* was higher in immatures compared to adults, while immatures also had higher back *L*, *a*, and *b* measurements (Fig. 3, Table 2). Together, these results confirm that immature birds included in our study tended to have lighter, more ‘buffy’ backs compared to adult juncos.

**Figure 3:**
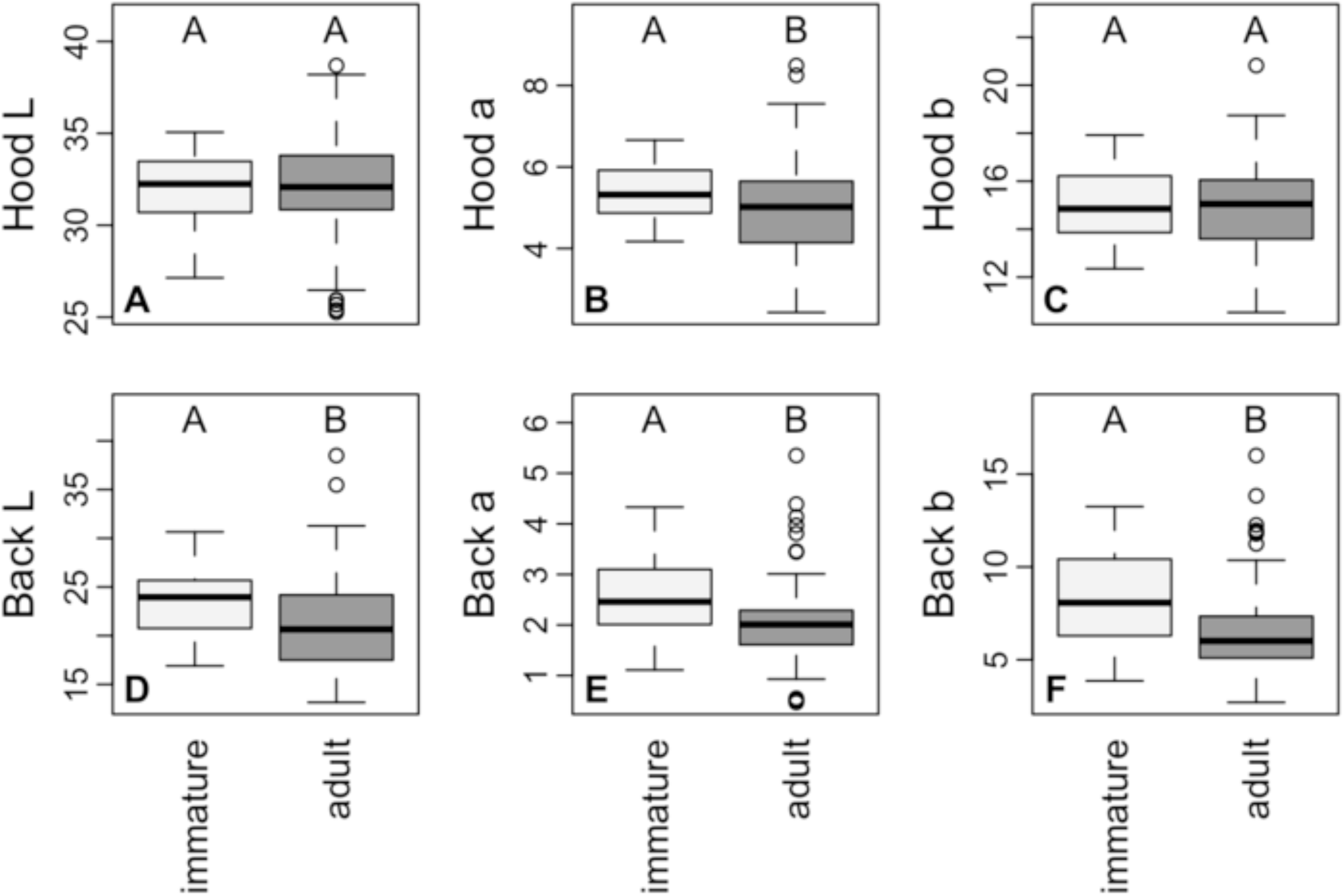
Boxplots of hood and back coloration measurements among immature and adult individuals of the “Oregon” Dark-eyed Junco complex. For the back and hood, “L” corresponds to brightness with higher values indicating brighter colors; “a” corresponds to redness, in which higher values correspond to more red coloration; and “b” corresponds to yellowness, in which higher values correspond to more yellow coloration. Shown above each box is the group classification following a Tukey’s HSD test for pairwise differences in mean values with the alphabetical order of groupings corresponding to descending differences in mean values among groups.

We also documented differences in coloration among subspecies of the “Oregon” Dark-eyed Junco group (Fig. 4, Table 2). We observed greater differentiation among subspecies in the mean values of hood measurements than back measurements. Using the Tukey’s HSD test, we identified as many as four groups of mean hood coloration values among subspecies (Fig. 4A, Fig. 4B, Fig 4C). The northernmost subspecies, *J. h. oreganus*, had the lowest average hood reflectance, or *L* values, while *J. h. Thurberi* had the highest average *L* values. The resident subspecies, *J. h. pinosus*, had the highest average hood *a* values, or ‘redness’, while *J. h. montanus* had the lowest average hood *a* values. Patterns for hood *b* values, or ‘yellowness’, followed a similar pattern to hood *a* values, but exhibited more overlap among subspecies.

**Figure 4:**
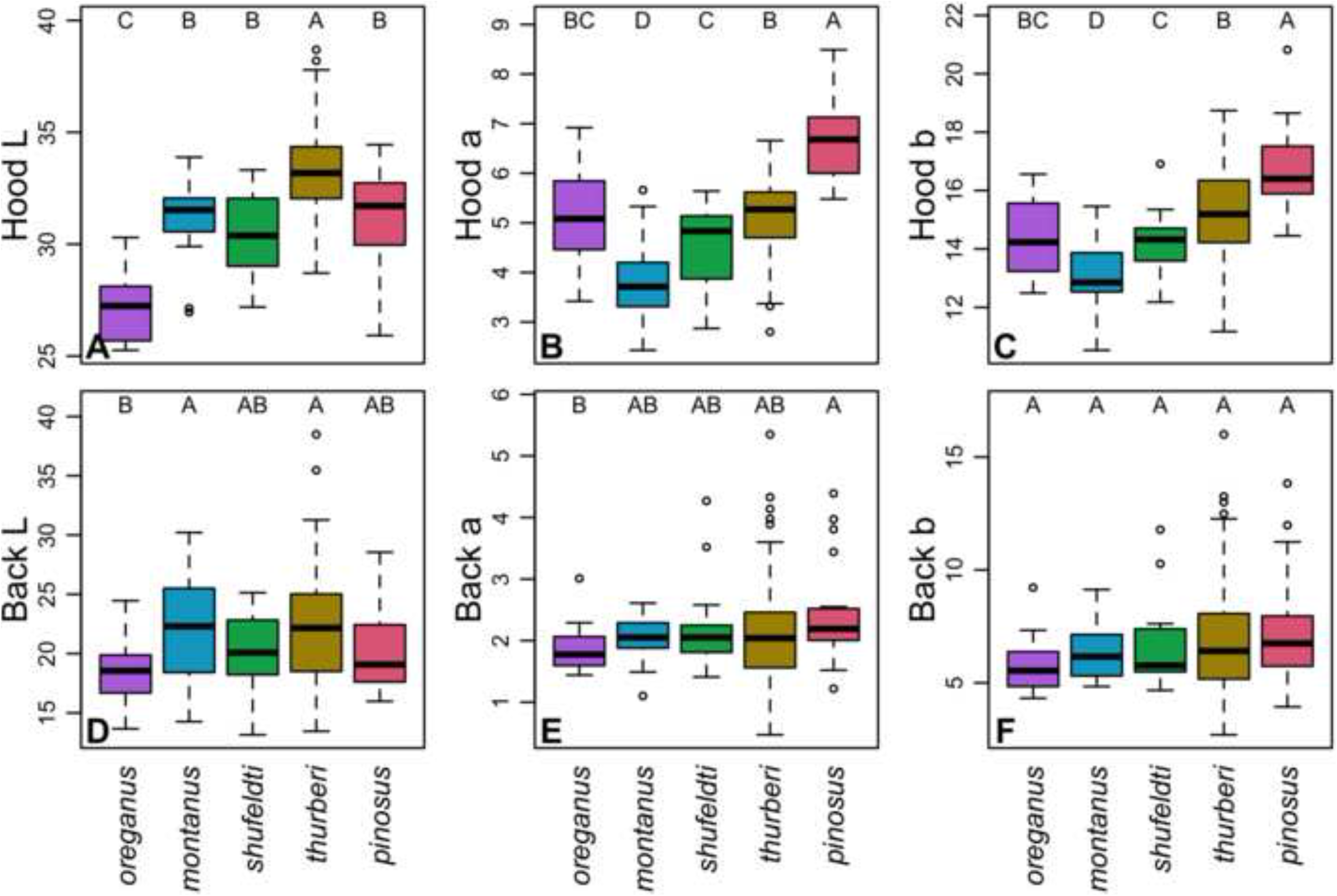
Boxplots of hood and back coloration among subspecies of the “Oregon” Dark-eyed Junco complex. For the back and hood, “L” corresponds to brightness with higher values indicating brighter colors; “a” corresponds to redness, in which higher values correspond to more red coloration; and “b” corresponds to yellowness, in which higher values correspond to more yellow coloration. Shown above each box is the group classification following a Tukey’s HSD test for pairwise differences in mean values with the alphabetical order of groupings corresponding to descending differences in mean values among groups.

**Figure 5:**
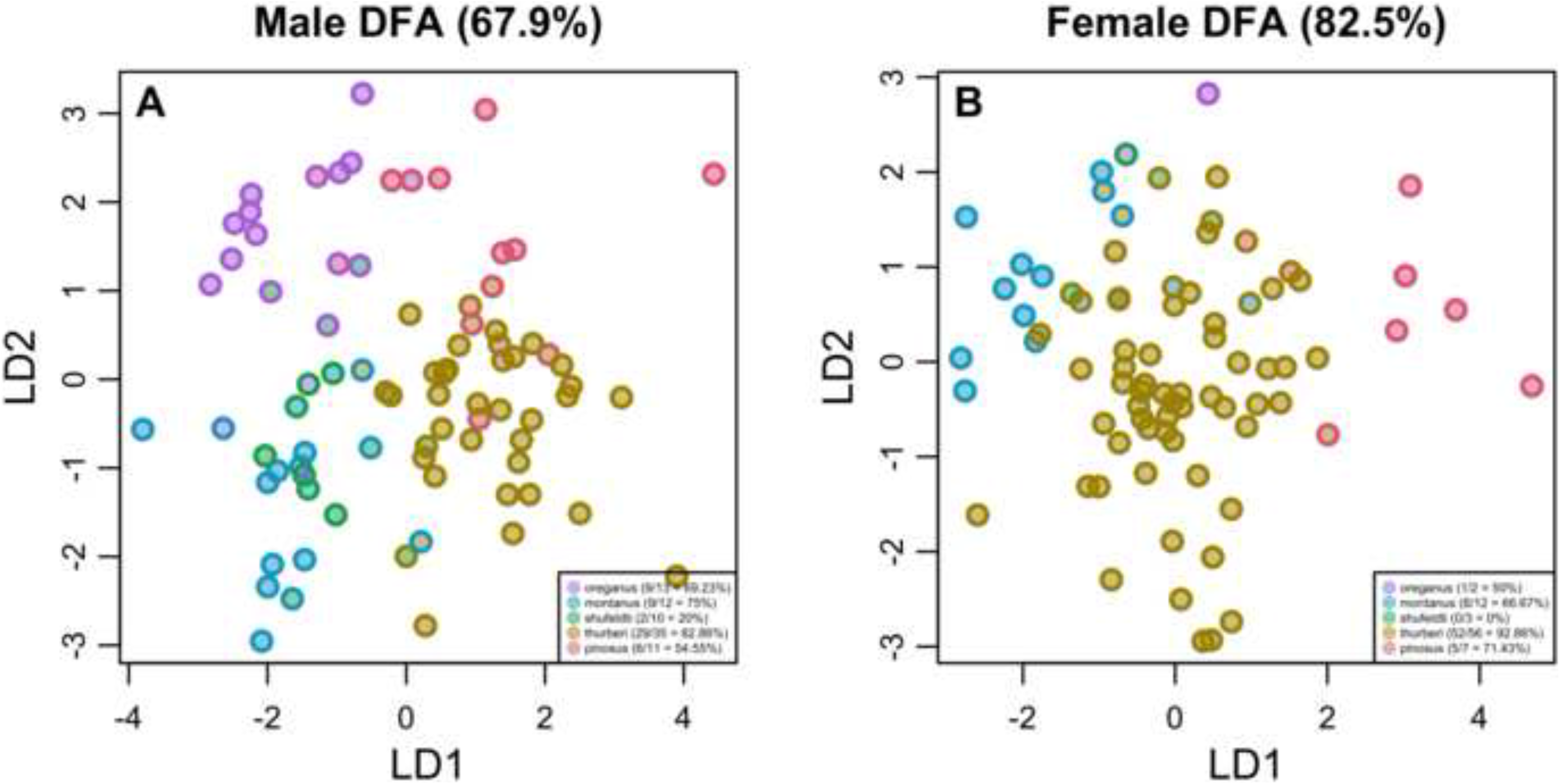
Discriminant function analysis and cross-validation analyses based on hood and back coloration for adult (A) males and (B) females among subspecies within the “Oregon” Dark-eyed Junco complex. The percentage of total correct classifications across subspecies is shown at the top of each plot. Within each plot, the center of each point corresponds to the subspecies identity associated with the metadata of each specimen, while the outside ring of each point corresponds to the predicted subspecies of each individual.

Although average hood measurements varied substantially among subspecies, differences in back coloration were less pronounced (Fig. 4, Table 2). The maximum number of back groupings we recovered via Tukey’s HSD was two (Fig. 4D, Fig. 4E, Fig 4F). Differences among subspecies in average back *L* values, or reflectance, were similar to that observed among hoods, albeit with greater overlap among subspecies.

The number of days since molt did not exhibit any associations with any back or hood measurements. The number of years since collection exhibited positive correlations with hood *b*, back *a*, and back *b* values (Table 3), consistent with a pattern of ‘foxxing’, or increasing red and yellow with years collection among the specimens studied here.

**Table 3:** Results from diagnosability tests of the “75% rule” formalized by Patten and Unitt (2002) using the first two axes of the discriminant function analysis with all five subspecies included. When the test statistic (D) is greater than zero, then 75% of the distribution for the first subspecies lies outside of 99% of the other subspecies.

|  |  | Male |  | Female |  |
| --- | --- | --- | --- | --- | --- |
| Subspecies 1 | Subspecies 2 | D<sub>12</sub> | D<sub>21</sub> | D<sub>12</sub> | D<sub>21</sub> |
| *oreganus* | *montanus* | 0.00 | -0.61 | **1.68** | -66.93 |
| *oreganus* | *shufeldti* | -1.67 | -1.09 | -6.57 | -9.55 |
| *montanus* | *shufeldti* | -1.11 | -0.81 | -6.85 | -0.20 |
| *shufeldti* | *thurberi* | -0.71 | -0.86 | -1.02 | -1.99 |
| *thurberi* | *pinosus* | -1.04 | -0.75 | -0.98 | **0.06** |

The discriminant function analysis using the colorimetry data on adult specimens correctly assigned 55 out of 81 males to their identified subspecies (67.90%; Fig. 4A) and 66 out of 80 females (82.50%; Fig. 4B). Diagnosability varied among subspecies. For males, hood and back color successfully categorized (from north to south) 69.23% of *oreganus* (9/13), 75.00% of *montanus* (9/12), 20.00% of *shufeldti* (2/10), 82.86% of *thurberi* (29/35), and 54.55% of *pinosus* (6/11). For females, hood and back color correctly categorized 50.00% of *oreganus* (1/2), 66.66% of *montanus* (8/12), 0% of *shufeldti* (0/3), 92.86% of *thurberi* (52/56), and 71.43% of *pinosus* (5/7).

Among all pairs of subspecies and sexes for which we tested the “75% rule”, only two compairisons were diagnosable with *D*_*ij*_ values ≥ 0: female *J. h. oreganus* were diagnosable from female *J. h. montanus* (*D*_*ij*_ = 1.68; Table 3) and female *J. h. thurberi* were diagnosable from *J. h. pinosus* (*D*_*ij*_ = 0.06; Table 3). In both these cases, the converse comparisons did not meet the requirements of the “75% rule”. All other pairwise comparisons had largely overlapping distributions of discriminant function scores and therefore failed to meet the “75% rule” threshold of diagnosability.

## Discussion

Using a colorimeter, we documented variation in the coloration of hoods and backs between subspecies, sexes, and age classes within the “Oregon” Dark-eyed Junco complex (Table 2; Fig. 2; Fig. 3; Fig. 4). Diagnosability of Oregon Junco subspecies on the basis of back and hood coloration was limited for both sexes. Only two subspecies pairwise comparisons — male *oreganus* and *montanus* and female *oreganus* and *montanus* — passed the ‘75% rule’ often used to delimit intraspecific taxonomy in birds (Table 2; Patten & Unitt 2002). Furthermore, cross-validation of our DFA was only able to accurately predict the subspecies grouping of 67.90% of males and 82.50% of females (Figure 4). In contrast, Miller (1941) claimed that back color allows a 90% “separation rate” between *shufeldti* and *thurberi* in interior ranges and a 75% separation rate along the coast, a 92% separation rate between *oreganus* and *shufeldti*, and a 97% separation between *oreganus* and *montanus*. Similarly high separation rates are reported for other subspecies pairs. When comparing *montanus* and *shufeldti*, Miller (1941) reported a 75% to 80% separation rate, but acknowledged that *shufeldti* is variable enough to “include the original *montanus* series.” It is unclear the methodology Miller (1941) used when generating this output and whether the numbers represent the extent of overlap, correct identification rate based on back color, or another method of differentiation. The most direct interpretation is that they represent the percentage of species that were able to be identified to subspecies by back color alone. In this case, our diagnosability rates fell short of what Miller (1941) reported, suggesting that subspecies of Oregon Dark-eyed Juncos exhibit weaker differentiation in dorsal coloration than has been heretofore assumed. Importantly, we had low sample sizes for females in a few subspecies, most notably the non-migratory *pinosus* (n_male_ = 11, n_female_ = 7), which has a restricted geographic range and correspondingly few specimens in most collections. We also had low sample sizes for *oreganus* (n_male_ = 13, n_female_ = 2), which decreased our statistical power to detect diagnosable differences for these taxa and contributed to the output of the “75% rule” tests involving *oreganus* and *pinosus*.

Beyond dorsal coloration, many subspecies in the Oregon Dark-eyed Junco complex exhibit broadly overlapping, clinal variation in other phenotypes. For example, many subspecies exhibit substantial overlap in morphological characters, such as tail length, wing length, and the extent of white on rectrices (Ferree 2013). Furthermore, four out of five of the Oregon Dark-eyed Junco subspecies included in this study (*thurberi*, *shufeldti*, *montanus*, and *oreganus*) exhibit little to no genetic population structure based on genomic analyses involving thousands of loci (Friis et al. 2018). *Pinosus*, however, exhibits pronounced genomic differentiation (Friis et al. 2018), is non-migratory, and has the shortest wings of all Oregon Dark-eyed Junco subspecies (Ferree 2013), and is therefore a valid subspecies. Although *oreganus* exhibits less genomic differentiation than *pinosus*, *oreganus* is also distinct in its darker plumage and partially geographically isolated range, suggesting it too may be a valid subspecies. On the other hand, *montanus*, *shufeldti*, and *thurberi* are less distinct: while *montanus* is duller than either *shufeldti* or *thurberi*, the broadly overlapping ranges between *shufeldti* with both *thurberi* and *montanus* and low genomic differentiation suggest a potential taxonomic revision for the three subspecies. Based on observed similarity in phenotype and genotype, one taxonomic solution may be to treat *montanus, shufeldti, and thurberi* as a single, widely distributed taxon with broad clinal variation across its range. Another solution would be to either lump *shufeldti* and *thurberi* together or to lump *thurberi* and *montanus*. Finally, a fourth option would be to retain all existing taxonomic classifications and continue to recognize *montanus, shufeldti*, and *thurberi* as distinct subspecies.

In summary, our quantitative, colorimetric analysis of dorsal values do not support preexisting assertions of subspecies diagnosability within the Oregon junco complex. Miller (1941)’s method for assessing pigmentary characters, which consisted of matching color effect in a given area of plumage with a graded series of color swatches and microscopic examination of pheomelanin and eumelanin, yielded results that are inconsistent with our colorimetry data. Our findings suggest a possible taxonomic revision of the Oregon Dark-eyed Junco group whereby fewer subspecies are recognized in light of broadly overlapping phenotypic and genetic variation. Revising subspecies limits in light of an improved understanding of geographic variation among populations promotes a more accurate and functional taxonomic classification of birds, which has broad implications across ornithology.

## Acknowledgements

We thank Philip Unitt and staff at the San Diego Natural History Museum, Kimball L. Garrett and staff at the Natural History Museum of Los Angeles County, and Kathy C. Molina and staff at the UCLA Donald R. Dickey Bird and Mammal Collection for providing access to specimens included in this study. We thank Zoe Yoo for their help in generating shape files used in the distribution map. NAM was supported by a NSF Postdoctoral Research Fellowship in Biology (1710739).

